# SCAND1 suppresses *CDC37* gene transcription by repressing MZF1

**DOI:** 10.1101/617753

**Authors:** Takanori Eguchi, Thomas L. Prince, Manh Tien Tran, Chiharu Sogawa, Benjamin J. Lang, Stuart K. Calderwood

## Abstract

CDC37 increases the stability of HSP90 client proteins and is essential for numerous intracellular oncogenic signaling pathways. Elevated expression of CDC37 was found in prostate cancer cells, although the regulatory mechanisms through which CDC37 expression becomes increased are unknown. Here we show both positive and negative regulation of *CDC37* gene transcription by two members of the SCAN transcription factor family- MZF1 and SCAND1, respectively. Consensus DNA-binding motifs for MZF1 were abundant in the *CDC37* promoter region. MZF1 became bound to these regulatory sites and *trans*-activated the *CDC37* gene whereas MZF1 depletion decreased CDC37 transcrption and reduced tumorigenesis of prostate cancer cells. On the other hand, SCAND1, a zinc-fingerless SCAN box protein that potentially inhibits MZF1, accumulated at MZF1-binding sites in *CDC37* gene, negatively regulated *CDC37* gene and inhibited tumorigenesis. SCAND1 was abundantly expressed in normal prostate cells but was reduced in prostate cancer cells, suggesting a potential tumor suppressor role of SCAND1 in prostate cancer. These findings indicate that CDC37, a crucial protein in prostate cancer progression, is regulated reciprocally by MZF1 and SCAND1.

## Introduction

The cell division control 37 (CDC37) protein plays a fundamental role in chaperoning the protein kinase family and participates in cancer by maintaining the activity of protein kinases involved in cell proliferation and transformation. These include tyrosine kinases such as Src [1], and serine/threonine kinases in the Raf-ERK pathway [2], Akt [3], the inhibitor of NF-κB kinase (IKK) [4], and cyclin-dependent kinase 4 (CDK4) [5–7]. CDC37 functions primarily in a complex with heat shock protein 90 (HSP90) to mediate the 3-dimensional folding and structural integrity of client proteins kinases [1, 8, 9]. CDC37 is particularly significant in prostate cancer as its overexpression leads to prostate carcinogenesis in transgenic mice [10–12]. It has been suggested that the high levels of oncogenic proteins present in most cancers make them dependent on molecular chaperones, a state referred to as “chaperone addiction” [13, 14]. Thus, because of their large protein clienteles, the CDC37-HSP90 axis offers a critical target for inactivating multiple oncogenic pathways [13, 14]. Consequently, the inhibition of HSP90 in cancer is currently a major area of research [14, 15]. However, less is known regarding the regulation of CDC37 expression in cancer and we have addressed this deficiency in this study.

Examination of the *CDC37* 5’ upstream region and introns indicated multiple consensus sequences that are potentially bound by the transcription factor Myeloid Zinc Finger 1 (MZF1, a.k.a. ZSCAN6, ZNF42, ZFP98). We recently reported that frequent amplification of MZF1 was observed in human cancers, further suggesting an oncogenic role for this factor [16]. Early studies showed that MZF1 might play roles in stemness and in the differentiation of hematopoietic stem cells and indicated its involvement in myeloma [17]. More recently, MZF1 was shown to participate in the malignant properties of several major solid tumors, including breast [18, 19], lung [20], liver [21], colorectal, and cervical cancer [22]. MZF1 protein structure is composed of an N-terminal SCAN domain, a linker region, and a C-terminal DNA binding domain that contains 13 repeats of zinc finger (ZF) motifs and is a member of the SCAN zinc finger (SCAN-ZF) family [16, 23]. The SCAN domain is leucine-rich oligomerization domain and is highly conserved in the SCAN-ZF family, which is composed of more than 50 family members [24–26]. Zinc fingerless SCAN domain-only proteins also exist [23]. SCAND1 is a SCAN domain-only protein that has been shown to bind MZF1 and other SCAN-ZF family members. SCAN domain-only proteins have been suspected to be suppressors of intact SCAN-ZF transcriptional activity [23, 24]. This possibility however has remained experimentally untested. Furthermore, we have tested the hypothesis that the relationship between MZF1 and SCAND1 regulates CDC37 expression and thereby cancer growth.

In this report we have, for the first time, characterized the *CDC37* gene promoter and determined that MZF1 indeed increases *CDC37* expression, while SCAND1 represses CDC37 expression. Our findings provide insight into how MZF1-driven *CDC37* expression promotes cancer progression and how SCAND1 functions as a potential tumor suppressor by repressing *CDC37*.

## Results

### Elevated expression of CDC37 is caused by MZF1 in prostate cancer

We first compared the expression levels of CDC37 between prostate cancer cell lines DU-145 and LNCaP, a castration-resistant prostate cancer (CRPC) cell line PC-3, and normal prostate cell line RWPE-1. CDC37 levels in the prostate cancer cells (PC-3, LNCaP, and DU-145) were higher than that of the normal cell line (Fig 1A). CDC37 levels in PC-3 cells were higher than that of the other prostate cancer cells in both confluent and growing conditions (Fig 1A).

**Fig 1.**
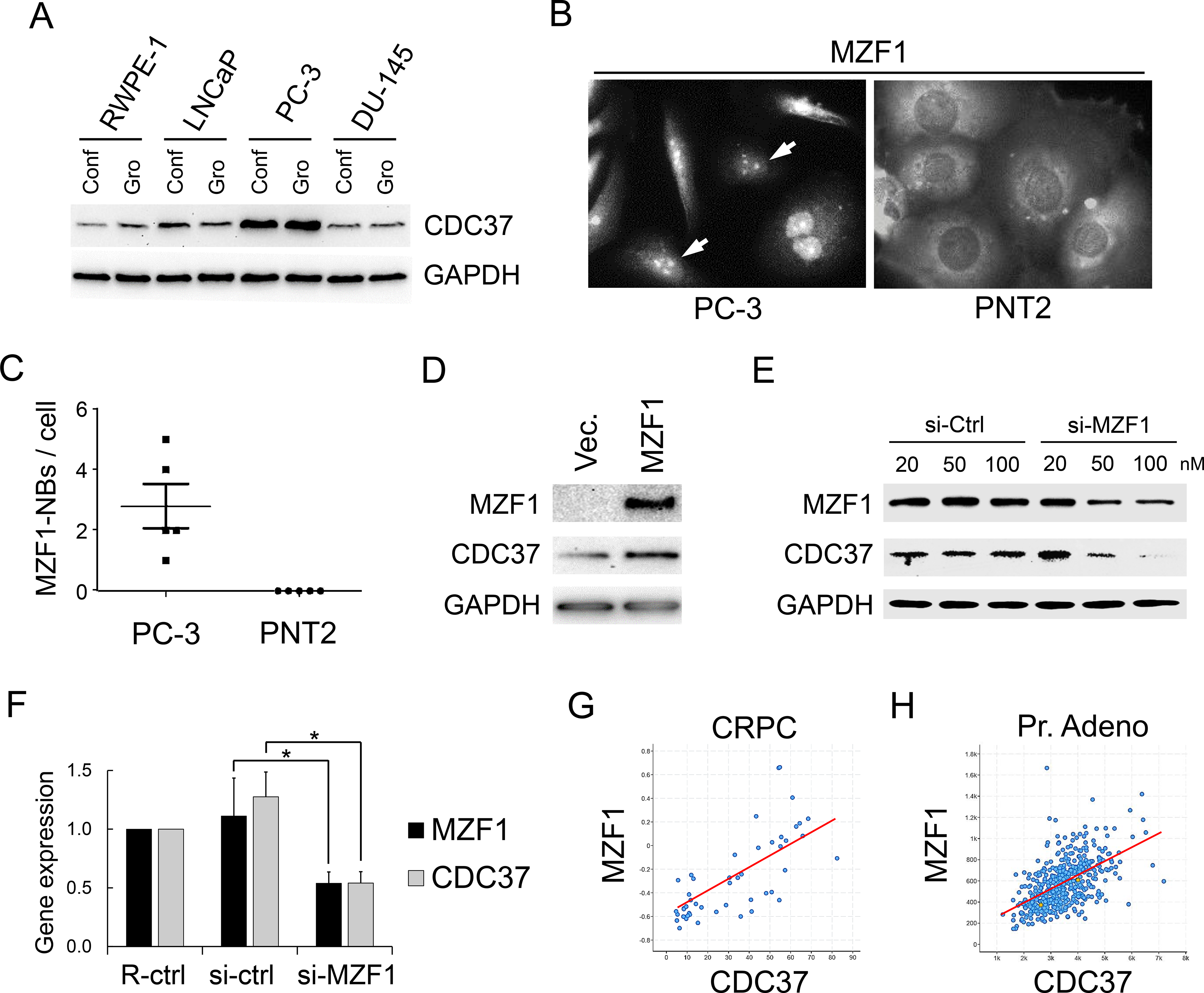
Elevated expression of CDC37 is caused by MZF1 in prostate cancer. Western blot showing CDC37 in prostate cancer cells. Cell lysates were prepared from confluent (Conf) and growing (Gro) RWPE-1, LNCaP, PC-3, and DU-145 cells. (B) Immunocytochemistry showing MZF1 expression and localization in PC-3 and PNT2 cells. Arrows indicate nuclear bodies of MZF1 (MZF1-NBs) seen in PC-3 but not in PNT2. (C) The number of MZF1-NBs per cells. (D) Western blot showing CDC37 altered by overexpressed MZF1 in DU-145. (E) Western blot showing CDC37 altered by depletion of MZF1 in DU-145 cells. (F) mRNA levels of CDC37 and MZF1 altered by depletion of MZF1. The MZF1-targeting siRNA mixture (A+B+C, 10 nM each) or a non-silencing control (si-ctrl) was transfected into PC-3 cells. RPL32, internal control. R-ctrl, reagent only control. *p < 0.05, n = 3. Similar results were obtained from 3 independent experiments. (G) Co-expression of MZF1 and CDC37 in patient samples of castration resistant prostate cancer (CRPC). N=114, Spearman correlation score 0.78, P=2.60e-11. (H) Co-expression of MZF1 and CDC37 in patient samples of prostate adenocarcinoma (Pr. Adeno). N=494, Spearman correlation score 0.57, P=7.74e-44

Although there are several possible mechanisms for the elevation in CDC37 expression in prostate cancer, in the present study we have focused on transcriptional regulation as the most immediate level of control. To determine potential *cis*-acting regulatory elements in the gene, we analyzed the DNA sequence around *CDC37* gene, between the −3.6kbp position and +2kbp position counted from the transcription start site (TSS) and found numerous consensus MZF1 binding sequences (Fig S1).

To query increased CDC37 level in PC-3 shown in Fig 1A, we next examined MZF1 expression and localization. MZF1 localized in nuclei in PC-3 but in the cytoplasm in PNT2 (Fig 1B), suggesting that both expression and nuclear localization of MZF1 could be involved in the increased CDC37 level in PC-3 cells. Additionally, nuclear bodies of MZF1 (designated MZF1-NBs) were seen in PC-3 but not in prostatic normal PNT2 cells, although the significance of the MZF1-NB is currently not known (Fig 1B, C). Moreover, MZF1 overexpression led to increases in CDC37 levels in PC-3 cells (Fig 1D) while siRNA-mediated knockdown of MZF1 lowered CDC37 mRNA and protein levels (Fig 1E, F). A role for MZF1 in CDC37 transcription was thus suspected.

To investigate the potential clinical significance of MZF1 regulation of CDC37, we next examined co-expression correlation of MZF1 and CDC37 in prostate cancer patient-derived tumor samples. Co-expression correlation was found between MZF1 and CDC37 in CRPC patient samples (Spearman’s rank correlation score: 0.78) and in prostate adenocarcinoma patient samples (Spearman’s correlation: 0.41). We next examined MZF1 and CDC37 expression correlation with prognosis of prostate cancer patients. High expression of MZF1 (P=0.0287) and CDC37 (P=0.0182) were correlated with poor prognosis of patients suffering from prostate cancer (Table 1).

**Table 1.** Correlation between gene expression and prognosis of prostate cancer patients.

|  | P value | N (total) | N (high) | N (low) |
| --- | --- | --- | --- | --- |
| *MZF1* | **0.0287** | 494 | 99 | 395 |
| *CDC37* | **0.0182** | 494 | 198 | 296 |
| *PSA* | 0.154 | 494 | 389 | 105 |

These data indicated MZF1 to be a potentially causal transcription factor for elevated expression of CDC37 in prostate cancer.

### SCAN zinc finger protein MZF1 directly *trans*-activates the *CDC37* gene

We next queried whether the SCAN zinc finger MZF1 could directly *trans*-regulate the *CDC37* gene through direct binding to *cis*-elements (MZF1 binding sequences) abundantly found in the *CDC37* 5’-upstream region. We examined the activities of *CDC37* promoter-driven luciferase reporter constructs containing different deletion mutants of the 5’ region and examined potential trans-regulation by MZF1 and its SCANless truncation constructs of the factor (Fig 2A, B). The full-length native MZF1 markedly increased *CDC37* promoter activities (of 3.6k, 950, 500/utr, 500, 202/utr) (Fig 1C). The SCANless mutant slightly increased *CDC37* promoter activities (of 500/utr, 500, 202/utr) but the transcriptional activity being much reduced compared with SCAN-containing native MZF1. These data indicated that SCAN oligomerization domain is essential for transcriptional activity of MZF1. It is notable that the 202/utr reporter was more potent in trans-activation compared to other constructs, suggesting the presence of potential inhibitory motifs upstream of the 202 sequence. We next examined whether the overexpressed MZF1 (with Flag-tag) could directly bind to *cis*-elements in genomic *CDC37* promoter region. MZF1 was detected by immunoblot in the crosslinked chromatin fraction and potentially post-translational modification forms (upshift), and potential trimer were detected in the chromatin (Fig 2D). The overexpressed MZF1 occupied genomic *CDC37* promoter regions (−0.4k and −1.8k regions that contain MZF1-binding sites) further indicating that the *CDC37* gene is regulated by MZF1 binding to these MZF1 binding sites in prostate cancer (Fig 2E).

**Fig 2.**
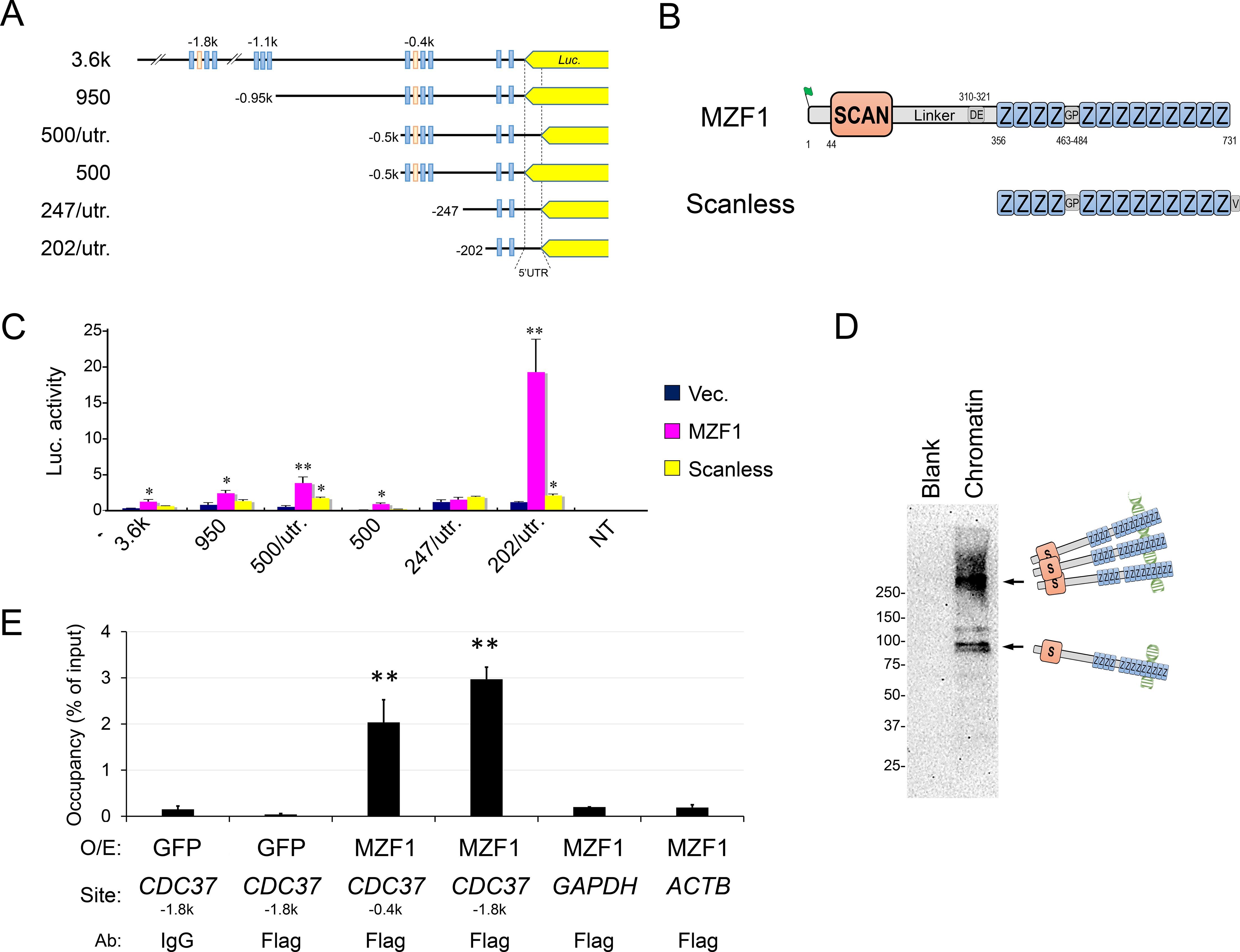
SCAN zinc finger MZF1 directly trans-activates *CDC37* gene. (A) Schemes of the *CDC37* promoter-reporter constructs. Truncated mutants of CDC37 promoter were connected with luciferase (*Luc*) gene. Blue box, MZF1 binding site. Orange box, heat shock element (HSE). 5’UTR, 5’ untranslated region. (B) The secondary structures of native MZF1 and the SCANless (zinc-finger domain alone). S, SCAN box. Z, zinc-finger motif. DE, aspartic acid- and glutamic acid-rich region. GP, glycine- and a proline-rich region. MZF1 was overexpressed with an N-terminal Flag-tag. The SCANless was overexpressed with a C-terminal V5-tag. (C) Luciferase activities from the truncated CDC37 promoters controlled by MZF1 and SCANless. Plasmid constructs shown in panels A and B were co-transfected into DU-145 cells. Vec., empty vector control. n=3, *P<0.05, **P<0.01 (vs Vec.). (D) Chromatin western blotting of MZF1 overexpressed in DU-145. Crosslinked chromatin was prepared from DU-145 overexpressed with MZF1. (E) ChIP-qPCR analysis showing MZF1 occupancy of *CDC37*. Chromatin was prepared from DU-145 overexpressed with Flag-MZF1 or GFP (control) and immunoprecipitated using anti-Flag antibody beads or control IgG. Co-immunoprecipitated DNA was analyzed by qPCR for *CDC37* (−1.8k or −0.4k regions) or control regions in *GAPDH* or *ACTB*. N=3, **P<0.01 (vs IgG control). Similar results were obtained from 3 independent experiments

These data indicate that MZF1 directly *trans*-activates *CDC37* gene through direct binding to *CDC37* promoter region and the SCAN domain-mediated trimerization is essential for full transcriptional activity of MZF1.

### The Zinc-fingerless SCAND1 factor suppresses *CDC37* gene and tumorigenesis of prostate cancer

The SCAN-ZF family consists of more than 50 members and contains a few potentially inhibitory zinc fingerless members such as SCAND1. We hypothesized that SCAND1 could repress *CDC37* gene transcription through oligomerizing with SCAN-ZF proteins such as MZF1. Notably, SCAND1 is highly expressed in normal prostate compared to the other tissues (http://biogps.org/#goto=genereport&id=51282). We found SCAND1 to be expressed in normal prostate cells while its levels were reduced in prostate cancer PC-3 and DU-145 cells (Fig 3A), suggesting that SCAND1 expression declined along with prostate oncogenesis. Overexpression of SCAND1 led to lowered CDC37 levels in PC-3 cells (Fig. 3B). To determine whether SCAND1 and a SCAN-only construct derived from MZF1 could regulate *CDC37* promoter constructs, we next carried out co-transfection and reporter assays. SCAND1 and SCAN-only constructs (SCAN and SCAN+linker) derived from truncation of MZF1 significantly repressed *CDC37* promoter activities (1.3k/utr, 500/utr, 202/utr) in PC-3 cells (Fig 3E). We also confirmed that the native MZF1 protein activated the CDC37 promoter whereas the SCANless mutant had little effect.

**Fig 3.**
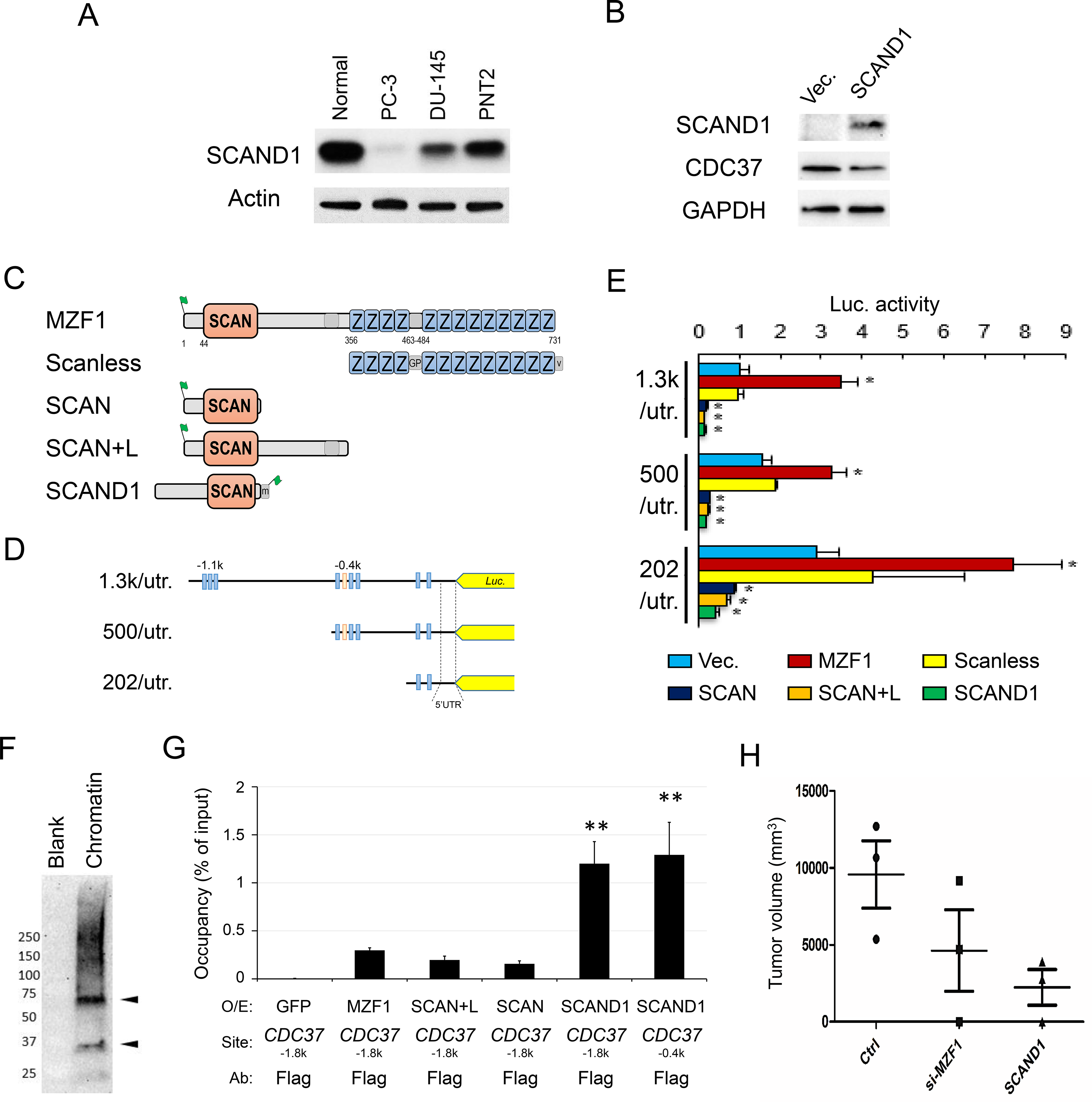
Zinc-fingerless SCAND1 suppresses *CDC37* gene and tumorigenesis of prostate cancer. (A) Western blot showing SCAND1 levels in normal prostate cells, PC-3, DU-145, and PNT2. (B) Western blot showing CDC37 lowered by SCAND1 overexpression. (C) Schemes of overexprssion constructs. (D) Schemes of truncated CDC37 promoters fused with luciferase reporter gene. (E) Luciferase activities from the truncated CDC37 promoters controlled by SCAND1 and truncated mutants of MZF1. Plasmid constructs shown in panels C and D were co-transfected into PC-3 cells. Vec., empty vector control. n=3, *P<0.05 (vs Vec.). (F) Chromatin western blot showing SCAND1 overexpressed in DU-145. Crosslinked chromatin was prepared from DU-145 overexpressed with SCAND1. (G) ChIP-qPCR analysis showing SCAND1 occupancy of CDC37. Chromatin was prepared from DU-145 overexpressed with SCAND1, MZF1, SCAN, SCAN+L or GFP (control) and immunoprecipitated using anti-Flag antibody beads. Co-immunoprecipitated DNA was analyzed by qPCR for CDC37 (−1.8k or −0.4k regions). N=3, **P<0.01 (vs IgG control). For figure 3 A-G, similar data were obatained from 3 independent experiments. (H) Tumorigenicity of PC-3 cells lowered by SCAND1 and depletion of MZF1. SCAND1-overexpression plasmid, MZF1-targeted siRNA, and control GFP plasmid were transfected into PC-3 cells which were then xenografted to SCID mice subcutaneously

We next hypothesized that SCAND1, although lacking ZF domains might be able to associate indirectly with MZF1-binding sites in genomic CDC37 promoter regions, perhaps through SCAND1-SCAN domain of MZF1 interaction. The overexpressed SCAND1 was detected in chromatin in PC-3 as monomer of the expected molecular weight. We also observed potential dimer and higher molecular weight forms in the crosslinked chromatin, suggesting higher oligomerization of SCAND1 on chromatin that might be consistent with a powerful repressor role (Fig 3F). The overexpressed SCAND1 (with Flag-tag) occupied the *CDC37* promoter region (−1.8k and −0.4k regions that we showed to contain MZF1-binding sites) (Fig 3G). The SCAND1 occupancy of the CDC37 regulatory site (−1.8k) was evidently much higher in magnitude than observed with MZF1 and SCAN-only constructs of MZF1 as suggested by increased enrichment in the ChIP assay (comparing Figs 2E, Fig 3G). These data indicated that SCAND1 could repress CDC37 gene powerfully through association with DNA sequences that we showed to bind MZF1. Therefore, our data indicate that while MZF1 positively regulates CDC37, SCAND1 negatively regulates transcription, a finding which could be crucial in mechanisms of prostate cancer progression.

We therefore hypothesized that, in this context, *MZF1* has an oncogenic influence while *SCAND1* may suppress tumorigenicity. To elucidate this possibility, we asked whether tumor initiation by prostate cancer PC-3 cells could be suppressed by SCAND1 expression and by depletion of MZF1. Indeed, SCAND1 suppressed tumorigenicity of PC-3 cells (Fig 3H). Depletion of MZF1 also lowered tumorigenicity of PC-3 cells.

## Discussion

Our data therefore suggest a novel mechanism for prostate cancer regulation by MZF1 (Fig. 1). CDC37 is a crucial molecule in prostate cancer growth through its fostering of oncogenic kinases and our data strongly indicate that the chaperone is regulated by MZF1. We show that MZF1 binds to sites in the CDC37 promoter and strongly activates transcription, and crucially that MZF1 expression is tightly correlated with CDC37 levels in clinical prostate cancer (Figs 1, 2). The ability of an endogenous inhibitor, the zinc fingerless SCAND1 factor to bind the promoter and repress transcription of MZF1 suggests a potential mechanism for prostate tumor suppression (Fig. 3). These data suggest mechanisms whereby members of the SCAN transcription factor family can fine-tune prostate cancer growth by up- or down-regulating CDC37 transcription and thus decide the outcome of prostate tumorigenesis (Fig 4). It is likely that other MZF1 targets important in tumorigenesis may be regulated in a similar way [16]. Our data do not exactly define the mechanism through which SCAND1 represses MZF1 activity although direct co-occupation of the *CDC37* promoter appears to be involved (Fig. 3). MZF1 was shown to bind DNA as a homodimer or heterodimer with other SCAN domain proteins and recruit chromatin remodeling protein mDomino [27]. Our data pinpoint the importance of the SCAN domain, as the SCANless construct of MZF1 showed much reduced transcriptional activity compared to the powerful activity of the native MZF1 (Fig. 3E). The SCAN domain is leucine-rich and mediates oligomer formation [16]. We found that MZF1 could form a higher order structure, potentially a trimer on chromatin (Fig. 2D). Therefore, it was indicated that SCAN domain of MZF1 is required for its oligomerization and thus for full activity for *trans*-activation. Little is currently known regarding the mechanisms whereby SCAND1 could repress transcription, although our data indicate that this may associate with the CDC37 promoter in oligomeric form at the site of MZF1 binding (Fig. 3).

**Fig 4.**
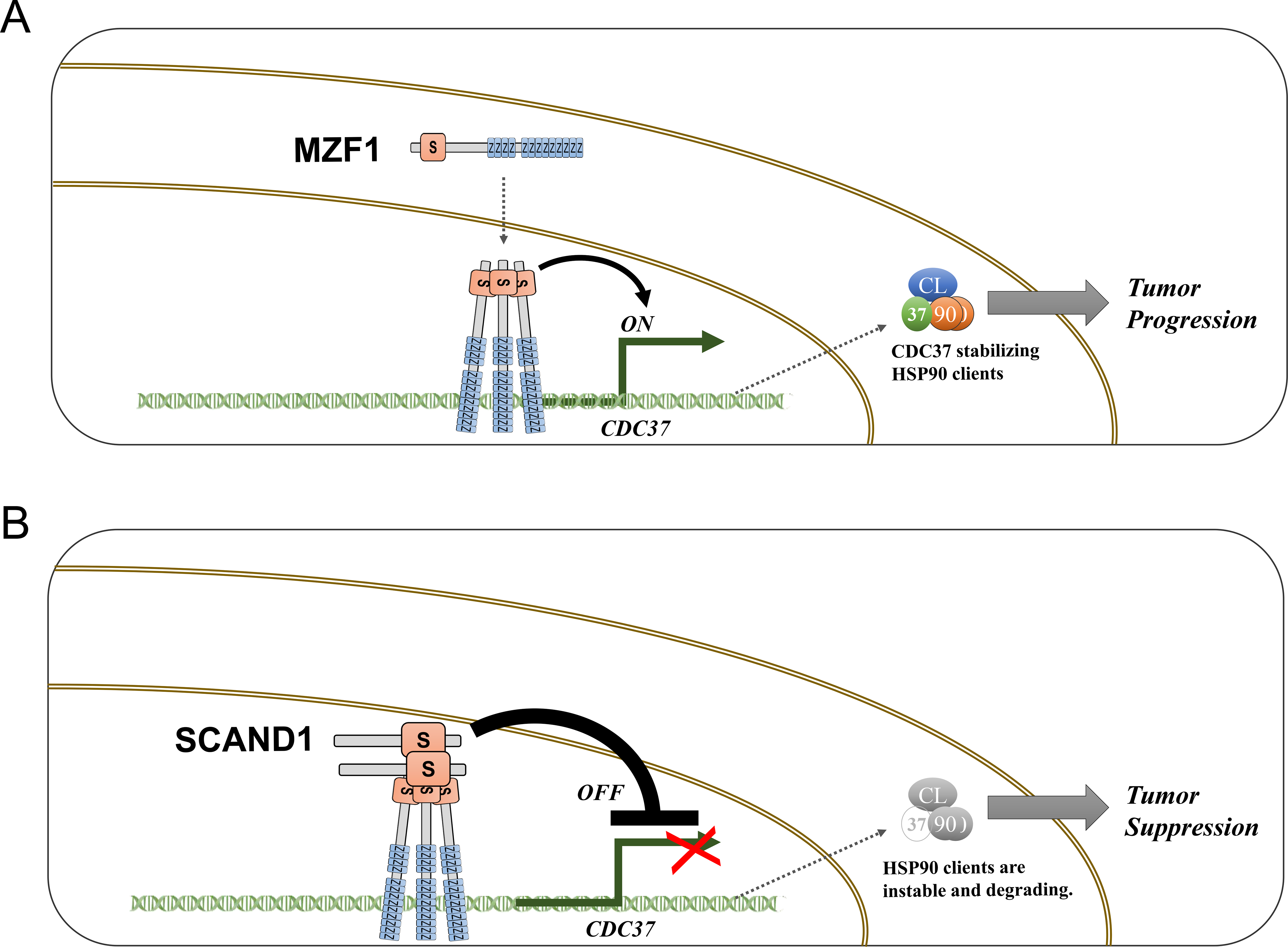
Graphical abstract: positive and negative regulation of the CDC37 gene by MZF1 and SCAND1 respectively, in prostate cancer. CDC37 is a key protein that stabilizes HSP90 client kinases essential for cancer progression. Elevated expression of CDC37 in prostate cancer is caused by MZF1 transcriptional activation whereas SCAND1 negatively regulates the CDC37 gene and prostate cancer tumorigenesis. CL, client. 37, CDC37. 90, HSP90

Our studies on CDC37 are aimed ultimately at targeting this factor in cancer as an alternative to Hsp90. Numerous HSP90 inhibitors have been developed to target the abundant HSP90, which plays versatile oncogenic roles in many types of cancer [28]. However, it has been difficult so far to employ Hsp90 inhibitors at effective doses due to normal tissue toxicity. Depletion of CDC37 reduced prostate cancer cell growth and attenuated MEK-ERK signaling pathway and PI3K-Akt signaling pathway [11]. Moreover, overexpression of CDC37 triggered prostatic tumorigenesis [10]. Our studies therefore suggest that further study of the role of MZF1 and SCAND1 in CDC37 transcription may indicate potential mechanism that may be targeted for inhibition of expression of the chaperone.

In conclusion, it was demonstrated that pro-oncogenic MZF1 and its antagonist SCAND1 coordinately regulate expression of CDC37, a factor that is essential for prostate cancer tumorigenesis. The SCAN domain is required for full transcriptional activity of MZF1 whereas SCAN-only factor SCAND1 powerfully associates with chromatin and represses tumorigenesis of prostate cancer cells.

## Materials and Methods

### Cell culture

PC-3, DU-145, LNCaP, and RWPE-1 cells were obtained from ATCC. PC-3 was cultured in F-12K or RPMI 1640 medium containing 5% to 10% fetal bovine serum (FBS). DU-145 was cultured in DMEM containing 5 to 10% FBS. LNCaP was cultured in RPMI 1640 medium containing 5% to 10% FBS. RWPE-1 was cultured in Keratinocyte Serum Free Medium (Thermo Fisher Scientific, Waltham, MA) supplemented with bovine pituitary extract (BPE) and human recombinant epidermal growth factor (hEGF). PNT2 was obtained from Sigma and cultured in RPMI 1640 medium supplemented with 2 mM glutamine and 5% to 10% FBS. Normal human prostate epithelial cells were purchased from Lonza (Basel, Switzerland) and cultured in Prostate Epithelial Cell Basal Medium (Lonza) supplemented with BPE, hydrocortisone, hEGF, epinephrine, transferrin, insulin, retinoic acid, triiodothyronine, and GA-1000.

### Molecular Cloning

Genomic DNA was isolated from a buccal cell swab of a deidentified white male. The 3.6 kbp human *CDC37* promoter region was PCR amplified and subcloned into a pGL3-luciferase vector (Promega, Madison, WI) using *Kpn*I and *Bam*HI. Once the pGL3-3.6 kbp CDC37promo vector was made, the 1.3 kbp, 950 kbp, 500 bp, 247 bp and 202 bp fragments with and without the 116-bp 5’UTR were subcloned into pGL3 similar using the same restriction sites as above. Each entire promoter fragment clone was Sanger sequenced. The complete 3.6 kbp *CDC37* promoter sequence was aligned with human reference genome GRCh37.

Human MZF1 cDNA was subcloned from pOTB-MZF1 into pcDNA3-Flag vector via TOPO directional cloning (Thermo Fisher Scientific) and designated pcDNA3/Flag-MZF1. For SCAN-only constructs, UGA stop codons were generated in the pcDNA/Flag-MZF1 via Quickchange mutagenesis and designated pcDNA3/Flag-C125/SCAN and pcDNA3/Flag-C252/SCAN+L. For SCANless, the cDNA of MZF1 zinc-finger domain was amplified via PCR and subcloned into pcDNA/V5 vector using TOPO-directional cloning and designated pcDNA/MZF1-V5/N252 (SCANless). Human SCAND1 (NM_033630) ORF clone with Myc-DDK C-terminal tag (RC200079) was purchased (Origene, Rockville, MD) and designated pCMV6-ScanD1-myc-Flag. pCMV-EGFP (NEPA Gene) was used as an overexpression control.

### *In silico* analysis of promoters and gene bodies

Sequences of promoter regions and gene bodies of human *CDC37, HSP90AA1* and *HSP90AB1* were obtained from the Eukaryotic Promoter Database [29]. Binding sites for MZF1 were predicted using PROMO [30, 31].

#### Luciferase Assay

Transient transfection and luciferase assays were performed as previously described [32]. Cells were cultured in 96-well plates. A plasmid (25 ng reporter, 100 ng effector) was transfected with 0.4 μl FuGENE HD (Roche, Basel, Switzerland) per well at a cell confluence level of 50-70%. The medium was changed at 16-20 hours after transfection. At 40-48 hours after transfection, 70 μl of the medium was aspirated, then 30 μl of Bright-Glo reagent (Promega, Madison, WI) was added and mixed by pipetting. Cells were incubated for 5 minutes at 37°C. The lysate (40 μl) was transferred to a 96-well white plate for measurement of luminescence.

### siRNA

The siRNA were designed based on siRNA design method of JBioS (Japan Bio Services, Saitama). RNA duplex of 19-bp plus TT-3’ overhangs in each strand was synthesized by Nippon Gene (Tokyo, Japan). For electroporation-transfection, total 40 pmol siRNA was transfected to 1 to 5 × 10^5^ cells. For reagent-transfection, cells were transfected with siRNA at the final concentration of 20-100 nM. Non-targeting siRNA was purchased from Nippon Gene (Tokyo, Japan). The designed sequences of siRNA were listed in Supple Table.

### Transfection

For ChIP assay and tumorigenesis assay, electroporation-mediated transfection was performed as described previously [33, 34]. To optimize electroporation for each cell type, cells (1 × 10^5^ to 1 × 10^6^ cells), plasmid DNA (2 to 10 μg total) or siRNA (40 pmol total), and serum-free medium were mixed to 100 μl total in a green cuvette with 1-mm gap (NepaGene, Ichikawa, Tokyo) and set to NEPA21 Super Electroporator (NepaGene). Poring pulse was optimized between 100V and 300V for 2.5 or 5.0 msec pulse length twice with 50 msec interval between the pulses and 10% decay rate with + polarity, as shown in a supplemental figure. Transfer pulse condition was five pulses at 20V for 50 msec pulse length with 50 msec interval between the pulses and 40% decay rate with +/− polarity. After electroporation, cells were recovered in serum-contained media. PC-3 (5 × 10^5^ cells) was electroporation-transfected with 10 μg plasmid DNA or 40 pmol siRNA with poring pulse at 175V for 2.5 msec pulse length twice and then cultured for 5 days and 1 × 10^6^ cells were subcutaneously injected to each SCID mouse.

For overexpression-ChIP assay, DU-145 (5 × 10^6^ cells) was transfected with 15 μg of pcDNA/Flag-MZF1 or pCMV-GFP with poring pulse at 175 V for 5 msec pulse length twice, cultured for 4 days in a 15-cm dish, and then used for ChIP assay.

For western blot analysis with overexpression of MZF1 and SCAND1, PC-3 cells were transfected with pcDNA3.1(-), pcDNA/Flag-MZF1, and pCMV6-SCAND1-myc-Flag using FuGENE6 (Roche).

For qRT-PCR after MZF1 depletion, PC-3 cells were transfected with a mixture of MZF1-siRNA A, B, and C (SR305183, OriGene, Rockville, MD) or non-silencing siRNA (SR30004, OriGene) using Lipofectamine RNAi max (Thermo Fisher Scientific).

For depletion of MZF1 and western blotting, DU-145 cultured in 6-well plate was transfected with siRNA hMZF1-all-NM_003422-53 using lipofectamine RNAi Max. Medium was replaced with fresh one at 24 hours after the transfection. Cells were lysed at 48 hours after the transfection.

### Chromatin

For protein overexpression and chromatin immunoprecipitation assay, DU-145 cells (5 × 10^6^ cells) were electroporation-transfected with pCMV6-ScanD1-myc-Flag, pcDNA3/Flag-MZF1, pcDNA3/Flag-SCAN, pcDNA/Flag-SCAN+L, and pCMV-GFP and then cultured for 3 days in 150-mm dishes. ChIP was done using SimpleChIP enzymatic chromatin IP kit with magnetic beads (Cell Signaling Technology). For optimization, chromatin DNA was sheared with MNase (0, 0.25, 0.5, 0.75, and 1 μl per reaction) and ultra-sonication at high- and low-power using Ultrasonic Homogenizer Smurt NR-50M (Microtec Nition) and analyzed within 2%-agarose gel electrophoresis. After the optimization, chromatin DNA was sheared with MNase (0.75 μl per reaction) and ultra-sonication at high-power. Overexpressed proteins in the chromatin fraction was analyzed by western blotting using anti-MZF1 antibody (Assay Biotechnology) and anti-SCAND1 antibody (ab64828, Abcam, Cambridge, UK). Anti-Flag M2 magnetic beads (Sigma) were used for ChIP.

DNA was purified from the eluate and from 1% and 10% input chromatin using QIA-Quick DNA purification kit (Qiagen, Hilden, Germany) and eluted within 50 μl of ddH_2_O. The purified DNA (3-5 μl) was used for ChIP-qPCR. Amplification specificity and efficiency were confirmed with melting curve analysis, slope analysis, and agarose gel electrophoresis. Primers were listed in Supple Table.

### Tumorigenesis

The efforts of all animal experiments were made to minimize suffering. The studies were carried out in strict accordance with the recommendations in the Guide for the Care and Use of Laboratory Animals of the Japanese Pharmacological Society. The protocol was approved by the Animal Care and Use Committee, Okayama University (Permit Number: OKU-2016219). All animals were held under specific pathogen-free conditions. PC-3 cells (5 × 10^5^) were transfected with 40 pmol MZF1-targeted siRNA, 10 μg pCMV6-ScanD1-myc-Flag or 10 μg pCMV-EGFP via electroporation, cultured for 5 days, and then 2.5 × 10^6^ cells were subcutaneously injected on a back of a 6-7-years-old SCID mouse (CLEA Japan, Tokyo). Tumor volumes were measured at day 41 post-injection period. The major axis (a) and minor axis (b) of tumors were measured with a caliper. The tumors were deemed to be ellipsoid and the volumes were calculated with a formula as follows: a tumor volume (V) ≒ 4πab^2^/3.

### Western blotting

To compare CDC37 levels among several types of cells, cells were cultured to reach confluent in 6-cm dish or sparsely grown in 10-cm dish. For endogenous SCAND1, protein samples were collected when cells reached sub-confluent. At day 2 after medium replacement, cells were washed with PBS and lysed within CelLytic M (Sigma, St. Louis, MO) for 15 min with gentle shaking. Lysates were collected and centrifuged at 14,000 × *g* for 15 min. Protein concentration was measured using BCA protein assay (Thermo Fisher Scientific). Protein samples (10-40 μg: equal amount) were loaded to SDS-PAGE. Proteins were transferred to PVDF membrane with a semi-dry method. The membranes were blocked within 5% skimmed milk in TBST for 1 hour. The membranes were incubated with primary antibodies over night at 4°C and with secondary antibodies for 1 hour at RT. The membranes were washed 3 times with TBST for 15 min after each antibody reactions. Chemiluminescence was detected with ChemiDoc MP Imaging System (Bio-Rad).

For crosslinked chromatin western blotting, see “chromatin” section. For overexpression of MZF1 and SCAND1 in PC-3, cells were transfected with pcDNA3/Flag-MZF1 or pcDNA3(-) using FuGENE6 (Roche) and cultured for 24 hours. Cells were washed with ice-cold PBS twice and lysed within RIPA buffer supplemented with protease phosphatase inhibitor cocktail (Thermo Fisher Scientific) at 4°C for 15 min, and then collected. The lysed cells were homogenized 10 times using a 25-gauge needle attached to a 1-ml syringe and then incubated on ice for 30 min. The lysate was centrifuged at 12,000 × *g* for 20 min at 4°C to remove debris. The same amount of the protein samples (10-50 μg) were used for SDS-PAGE and semi-dry transfer.

For depletion of MZF1, DU-145 cultured in 6-well plate was transfected with siRNA hMZF1-all-NM_003422-53 using lipofectamine RNAi Max following manufacture’s instruction. Medium was replaced with fresh one at 24 hours after the transfection. Cells were collected using trypsin and counted at 48 hours after the transfection. Cell lysate was prepared using RIPA buffer as described above. The equal amount of lysate (25 μg) was loaded to each lane for SDS-PAGE (4-20% TGX gel, BioRad). The proteins were transferred to a PVDF membrane with wet-transfer method at 40V for 16 hours on ice with a transfer buffer (25 mM Tris, 192 mM Glycine, 10% methanol, and 0.05% SDS). The membrane was washed 3 times with TBST for 15 min and then blocked 1 hour within 5% skimmed milk. MZF1, CDC37, HSP90α, and HSP90β were detected by western blotting.

The antibodies used were anti-MZF1 (C10502, Assay Biotechnology, 1:500), anti-CDC37 (D11A3, Cell Signaling Technology, 1:1000), anti-FLAG (Clone M2, Sigma), anti-SCAND1 (ab64828, Abcam, Cambridge, UK), and HRP-conjugated anti-GAPDH antibody (Clone 5A12, Fujifilm Wako, 1:5000).

### Immunocytochemistry

Immunocytochemistry was performed as described [35]. Cells cultured in 4-well chamber slides were fixed with 4% (wt/vol) paraformaldehyde in PBS for 15 min. Cells were permeabilized with 0.2% Triton X-100 in PBS for 15 min. Cells were blocked in IHC/ICC blocking buffer high protein (eBioscience, San Diego, CA) for 10 min and then reacted with anti-MZF1 antibody (1:50, C10502; Assay Biotechnology, Fremont, CA) and AlexaFluor488 secondary antibody (Thermo Fisher Scientific, 1:1000) in the blocking buffer. Cells were washed with PBS for 5 min twice between the steps. Cells were mounted with ProLong Gold AntiFade Reagent (Thermo Fisher Scientific). Fluorescence images were captured using Axio Vision microscope equipped with an AxioCam MR3 (Zeiss, Oberkochen, Germany).

### RT-qPCR

RT-qPCR was performed as previously described [35, 36]. Total RNA was prepared using an RNeasy RNA purification system (Qiagen, Hilden, Germany) with *DNase* I treatment. cDNA was synthesized using an RT kit for qRT-PCR (Qiagen) with a mixture of oligo dT and random primers. The cDNA pool was diluted 5- to 20-fold. The cDNA standard with permissible slopes of PCR efficiencies was prepared by step dilution of the cDNA pool for relative quantification of mRNA levels. In 20 μl of qPCR mix, 0.25 μM of each primer, 4-10 μl of diluted cDNA, and 10 μl SYBR green 2x Master Mix (Applied Biosystems, Waltham, MA) was added and reacted at 95°C for 10 min, followed by 40 cycles at 95°C for 15 seconds and 60°C for 1 min. Dissociation curves with specific single-peaked PCR and proper amplification slopes for PCR efficiency were confirmed. LinRegPCR software was used for baseline fluorescence correction, and calculation of Cq values and PCR efficiency of amplicons [37]. Primer sequences are listed in Supple Table.

### Gene expression in clinical samples

Co-expression of MZF1 and CDC37 was analyzed in TCGA in cBioPortal. Data sets of NEPC/CRPC (Trento/Cornell/Broad 2016, 114 samples) and prostate adenocarcinomas (TCGA, PanCancer Atlas; 494 patients/samples) were analyzed and Spearman’s rank correlation coefficient of co-expression was shown on the figure.

For correlation of gene expression with patient prognosis, Kaplan-Meier survival analysis was performed using 494 prostate cancer patient samples in Human Protein Atlas Database.

#### Statistics

Data were expressed as the means ± SD unless otherwise specified. Comparisons of 2 were done with an unpaired Student’s *t*-test.

## Author Contributions

TE, TLP, and SKC conceptualized and designed the study. SKC and TE acquired funding. TLP and TE constructed materials. TE, TMT, CS, TLP, and BJL devised methodologies and performed experimentation. TE wrote the original manuscript. SKC, TLP and TE edited the manuscript. All authors reviewed the manuscript.

## Abbreviations

*CDC37*: cell division control 37
*CRPC*: castration-resistant prostate cancer
*EV*: extracellular vesicle
*HS*: heat shock
*HSE*: heat shock element
*HSF1*: heat shock factor 1
*HSP*: heat shock protein
*MZF1*: myeloid zinc finger 1
*NEPC*: neuroendocrine prostate cancer
*SCAN*: SREZBP-CTfin51-AW1-Number 18 cDNA

## Acknowledgments

This paper is dedicated to the memory of one of our mentors, Professor Ken-ichi Kozaki, who passed away on May 29, 2016. The authors thank Barbara Wegiel, Eva Csizmadia, Ayesha Murshid, Yuka Okusha, Kuniaki Okamoto, and Yasutomo Nasu for valuable, illuminating discussion and encouragement. The authors thank Sachin Doshi for technical assistance.

## Supplemental Items

**Fig S1.**
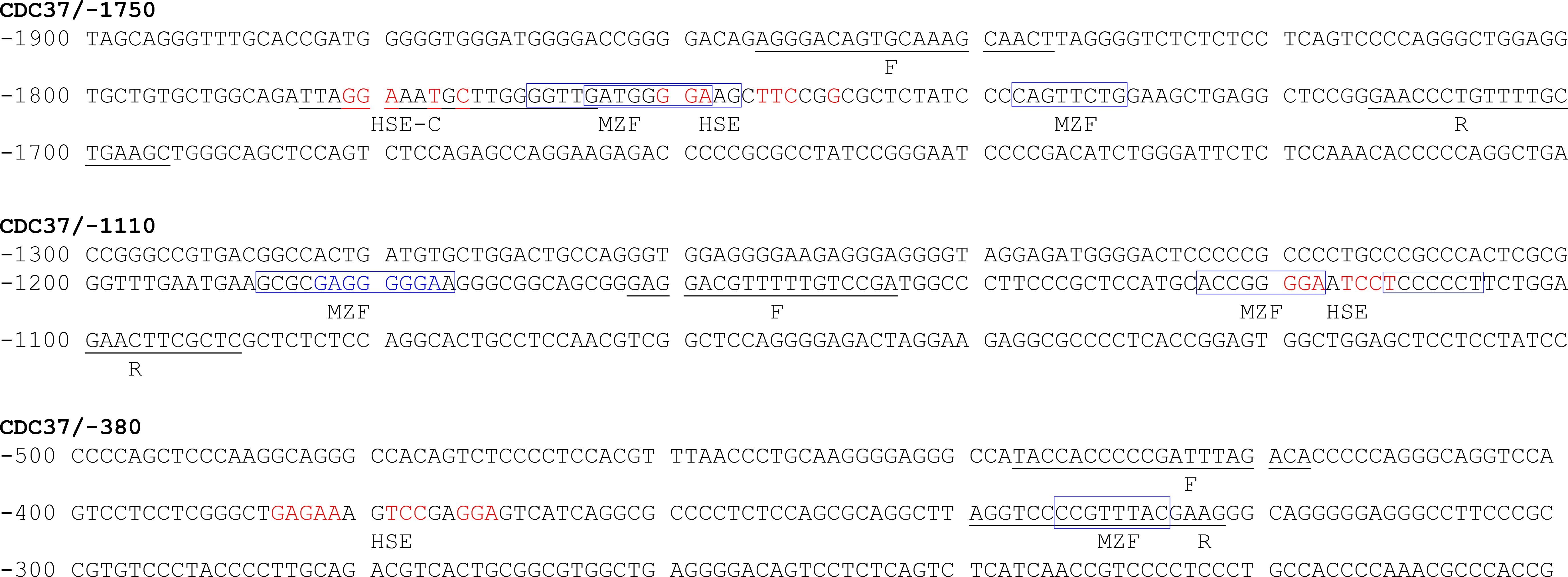
Binding sites for MZF1 and HSF1 in *CDC37*. MZF1 binding sites were enclosed with blue box. Heat shock elements (HSE) were shown with red. The sequences detected by forward primers (F) and reverse primers (R) in ChIP-qPCR were underlined

